# Ras-GRF1 in CRF Cells Controls the Early Adolescent Female Response to Repeated Stress

**DOI:** 10.1101/2020.02.19.955088

**Authors:** Shan-xue Jin, David A. Dickson, Jamie Maguire, Larry A. Feig

## Abstract

Ras-GRF1 (GRF1) is a calcium-stimulated guanine-nucleotide exchange factor that activates Ras and Rac GTPases. In hippocampal neurons, it mediates the action of NMDA and calcium-permeable AMPA glutamate receptors on specific forms of synaptic plasticity, learning, and memory in both male and female mice. Recently, we showed that GRF1 also regulates the HPA axis response to restraint stress, but only in female mice before puberty. In particular, we found that after exposure to 7-days of restraint-stress (7DRS) (30 min/day) elevation of serum CORT levels are suppressed in early adolescent (EA) female, but not EA male or adult female GRF1 knockdown mice. Here, we show that this phenotype is due, at least in part, to the loss of GRF1 expression in CRF cells of the paraventricular nucleus of the hypothalamus, as GRF1 knockdown specifically in these cells also reduces serum CORT response to 7DRS in EA females, but not EA males or adult females. Moreover, it reduces females CORT levels to those to found in comparably stressed control male mice. GRF1 knockdown in CRF cells also blocks the anxiolytic phenotype normally found in EA females 24 hrs after 7DRS. Interestingly, loss of GRF1 in these cells has no effect after only 3 exposures to restraint stress, revealing a role for GRF1 in repeated stress-induced CRF cell plasticity that appears to be specific to EA female mice. Overall, these findings indicate that GRF1 in CRF cells makes a key contribution to the distinct response early-adolescent female display to repeated stress.

## Background

The body’s reaction to environmental stressors involves sequential hormone release in the hypothalamus–pituitary–adrenal (HPA) axis (McEwen 2007). These perturbations stimulate corticosteroid (CORT) release from the adrenal cortex to mobilize an integrated protective response to stress. The first HPA axis response is the release of corticotropin releasing factor (CRF) from the terminals of parvocellular CRF hypothalamic neurons in the paraventricular nucleus (PVN) into hypophyseal portal vessels. Adrenocorticotropic hormone (ACTH) is then released from the pituitary, which stimulates corticosteroid synthesis and release from the adrenal gland.

Clear sex- and age-dependent differences in both how the HPA axis responds to stress, and how stress leads to psychiatric disorders have been documented (Bale and Epperson 2015). A consistent trend in animals demonstrates higher and longer-lasting CORT secretion after various stressors in adult females compared to males (Heck and Handa 2018). Sex differences at multiple sites in the HPA axis have been implicated, including pituitary response to ACTH, CRF expression, and glucocorticoid feedback (Bale and Epperson 2015; Heck and Handa 2018). Moreover, women are roughly twice as likely as men to suffer from stress associated disorders such as anxiety, PTSD, and depression (Bangasser and Valentino 2014).

Age-dependent differences in the response to stress have also been found. Adolescence is between ~10-25 yrs in humans and postnatal days 28-55 in rodents (Burke and Miczek 2014; Doremus-Fitzwater and Spear 2016). It can be divided into early, mid and late stages, with puberty beginning at mid-adolescence (Burke and Miczek 2014). Adolescence is a time of continuing brain maturation, particularly in regions that control the HPA axis (Juraska and Willing 2017). Excess stress during this crucial developmental stage may influence brain maturation and contribute to both anxiety and depression during adolescence, and psychiatric disorders later in life (Spear 2000). Consistent with this finding, many studies show that animals at various stages of adolescence respond differently to stress than adults. In general, it has been noted that exposure to a variety of stressors lead to greater or more prolonged hormonal responses in adolescents compared to adults (Romeo 2018). For example, prepubertal, early adolescent (EA) (25–30 days of age) male and female rats exposed to an acute stressor display a significantly extended hormonal response relative to adults (>65 days of age) (Romeo, et al. 2006). Moreover, after exposure to a combination of stressors, adolescent rats display reduced anxiety, rather than the enhanced anxiety typically seen in adults, that is associated with enhanced risk taking (Toledo-Rodriguez and Sandi 2011). In humans, the risk of depression reaches its maximum around early adolescence and decreases as humans age through mid and late-adolescence (McLaughlin and King 2015).

Sex differences in the stress response among adolescents have also been observed. For example, restraint stress significantly increased alcohol intake and preference in female adolescent rats, but decreased alcohol intake and preference in male adolescent and female adult rats (Wille-Bille, et al. 2017). Early adolescent (EA) specific differences between girls and boys have also been observed. For example, young girls exhibit greater CORT stress reactivity to social stress tests than boys (Blumenthal, et al. 2014; Blumenthal, et al. 2009; Gunnar, et al. 2009; McLaughlin and King 2015). Moreover, EA girls are more prone than EA boys to anxiety, depression and eating disorders, as well as suicidal ideation, while EA boys are more susceptible to ADHD (Hankin, et al. 1998; Skogli, et al. 2013).

PVN neurons that release CRF are tightly regulated in many ways. For example, with repeated exposure to stress, they display cellular, synaptic, and connectional plasticity that serves to maximize the ability of the HPA axis to maintain response flexibility (Aguilera and Liu 2012). At the cellular level, chronic stress often enhances the production of CRF and its co-secretagogue arginine vasopressin (AVP) and alters neurotransmitter receptor expression, so as to maximize cellular excitability. Interestingly, differences in the CRF cell response to stress have been found between male and female EA mice, such that social isolation, but not acute stress, alters the intrinsic properties of these cells in a sexually dimorphic fashion (Senst, et al. 2016).

p140-Ras-GRF1(GRF1) and p130-Ras-GRF2(GRF2) constitute a family of calmodulin-binding, calcium-activated exchange factors for both Ras and Rac GTPases that are expressed in neurons (not glia) throughout the adult central nervous system at similar levels in both males and females (Feig 2011). They have very similar domains including calmodulin (CaM)-binding IQ motifs, as well as Ras-activating CDC25 domains and Rac-activating DH domains. We showed that GRF1 mediates LTD induced by NR2B-type NMDA receptors through Rac/p38 Map kinase signaling (Li, et al. 2006), but high-frequency stimulated (HFS) LTP promoted by calcium permeable AMPA receptors (CP-AMPARs) via Rac/p38 signaling in the hippocampus (Jin, et al. 2013). GRF1 can also mediate dopamine signaling through Ras/Erk signaling in the striatum (Cerovic, et al. 2014). Recently, we showed using knockout mice that GRF1 also contributes to regulation of the HPA axis in mice, but in a sex, age, and stimulus dependent manner (Uzturk, et al. 2015). In particular, GRF1 contributes to the negative feedback the hippocampus plays in regulating the HPA axis response up to 5 exposures to 30min/day restraint stress, in prepuberal early-adolescent (EA) (pn-d28-35) females, but not their EA male, or late adolescent or adult female counterparts. However, we also showed, using these knockout mice, that GRF1 plays the opposite role after 7 exposures to restraint stress, where it contributes positively to the stress response specifically in EA females. It also contributes to the anxiolytic behavior we observed in EA female mice after restraint stress (Uzturk et al. 2015). Finally, puberty-associated hormonal changes do not contribute to this phenotype because the effects are observed before puberty and ovariectomy had no effect on GRF1 knockout phenotypes after puberty.

Here, by suppressing GRF1 function only in CRF cells in the PVN of the hypothalamus, we show that GRF1 is required in these cells for EA females to display their greater serum CORT response to 7 exposures of restraint stress (30 min/day) (7DRS) than that observed in EA males. Moreover, GRF1 is not required for a normal serum CORT response in EA male or adult female mice. Knockdown of GRF1 expression in CRF cells also prevented the 7DRS induced anxiolytic phenotype observed in EA female mice. In addition, we show that GRF1 in CRF cells of EA females does not influence the HPA response to fewer exposures to restraint stress, implicating GRF1 in a form of CRF cell plasticity likely unique to this cell type.

## RESULTS

### Inhibition of GRF1 expression in the PVN of the hypothalamus suppresses elevation of serum corticosterone levels induced by 7 days (30 min/day) of restraint stress specifically in early-adolescent (EA) female mice

We demonstrated previously that GRF1 negatively regulates the HPA axis response to short-term restraint stress (1-5 days; 30 min/day) in early-adolescent (EA) (prepubescent (peri-adolescent)) females (pn days 28-35)(Uzturk et al. 2015). In particular, GRF1 knockout mice display an exaggerated enhancement of serum corticosterone (CORT) levels after these exposures. No effect was detected in EA male, late adolescent (pn-days 35-42) female or adult female mice. This effect was found to be due to GRF1 in the CA1 hippocampus, consistent with the negative feedback role this brain region plays on the HPA axis.

However, we also showed that GRF1 plays the opposite role after 7 days of the same restraint stress, again specifically in EA females, where loss of GRF1 leads to a suppressed serum CORT response. We hypothesized that this difference is due to GRF1 function in a different brain area that plays a positive role in promoting the HPA axis to repeated stress. Thus, we first investigated the paraventricular nucleus (PVN) of the hypothalamus, because this region is known for this function and previous single cell RNAseq studies found that GRF1 is expressed in many cell types cells throughout the PVN, at least in adult males (Romanov, et al. 2017) (also see http://linnarssonlab.org/hypothalamus/). We expanded this finding, using GRF1-specific antibodies, as we detected positive staining of cells throughout the PVN of wild type EA female mice compared to results using GRF1 knockout mice (Fig. 1A and B).

**Figure. 1.**
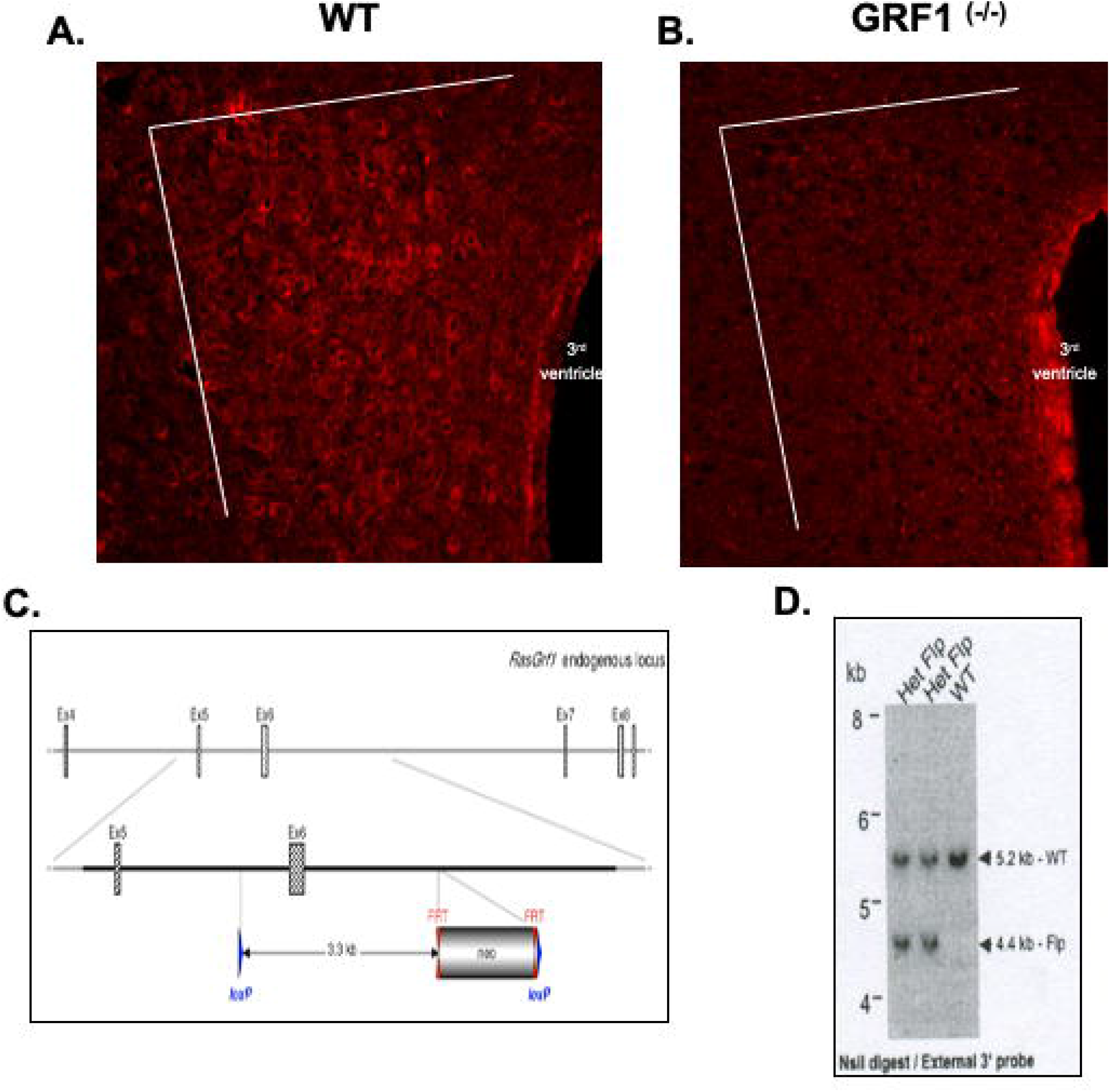
Generation of conditional GR1 knockout mice. **A-B.** GRF1 protein expression in the PVN of brain slices from WT mice (A) and whole animal GRF1^−/−^ knockout mice (B) stained with anti-rasGRF1 antibody (scale bar, 50 μm). White lines represent the borders of the PVN region. 3^rd^ ventricle can be visualized in the bottom right of both panels as labeled. **C.** Schematic representation of targeting strategy selected. Hatched rectangles represent RasGRF1 coding sequences, grey rectangles indicate non-coding exon portions, solid lines represent chromosome sequences. The neomycin positive selection cassette is indicated. loxP sites are represented by blue triangles and FRT sites by double red triangles. The size of the flanked RasGRF1 sequence to be deleted is specified. **D.** Genomic blots identifying heterozygous conditional knockout mice. Primer sequences available upon request.

Therefore, we tested whether GRF1 in the PVN hypothalamus contributes to stress-induced CORT production after repeated (7) daily episodes of restraint stress (30 min/day) (7DRS) in EA mice. To this end, the expression of GRF1 was inhibited in the PVN using stereotactic injection of adenovirus expressing CRE recombinase, or GFP as controls, into this brain region of 21-22-day old female floxed GRF1 mice (Fig. 1C and 1D). After waiting 7 days to allow virus expression, mice were exposed to 7DRS, or control handling, and then CORT levels in serum were measured. This timeline was employed so that at the end of the experiment mice were still in early-adolescence, because we showed that after this developmental stage, GRF1 no longer plays a role in HPA axis regulation (Uzturk et al. 2015). As expected, qPCR analysis of whole PVN RNA shows that injection of Adeno-CRE severely inhibited GRF1 mRNA levels compared to GFP controls in both EA females and males (Figure 2A). 2-way ANOVA (Factors: CRE, Sex) revealed a significant effect for injection of adeno-CRE (F_(1,8)_=428.3, P<0.0001) without a significant interaction between these terms (F_(1,8)_=0.2942, P=0.6023); Sidak *post-hoc* tests revealed a significant difference in GRF1 expression after injection of adeno-CRE in both females (t=15.70, *P*<0.0001) and males (t=13.85, *P*<0.0001) when compared to their adeno-GFP injected counterparts.

**Figure 2.**
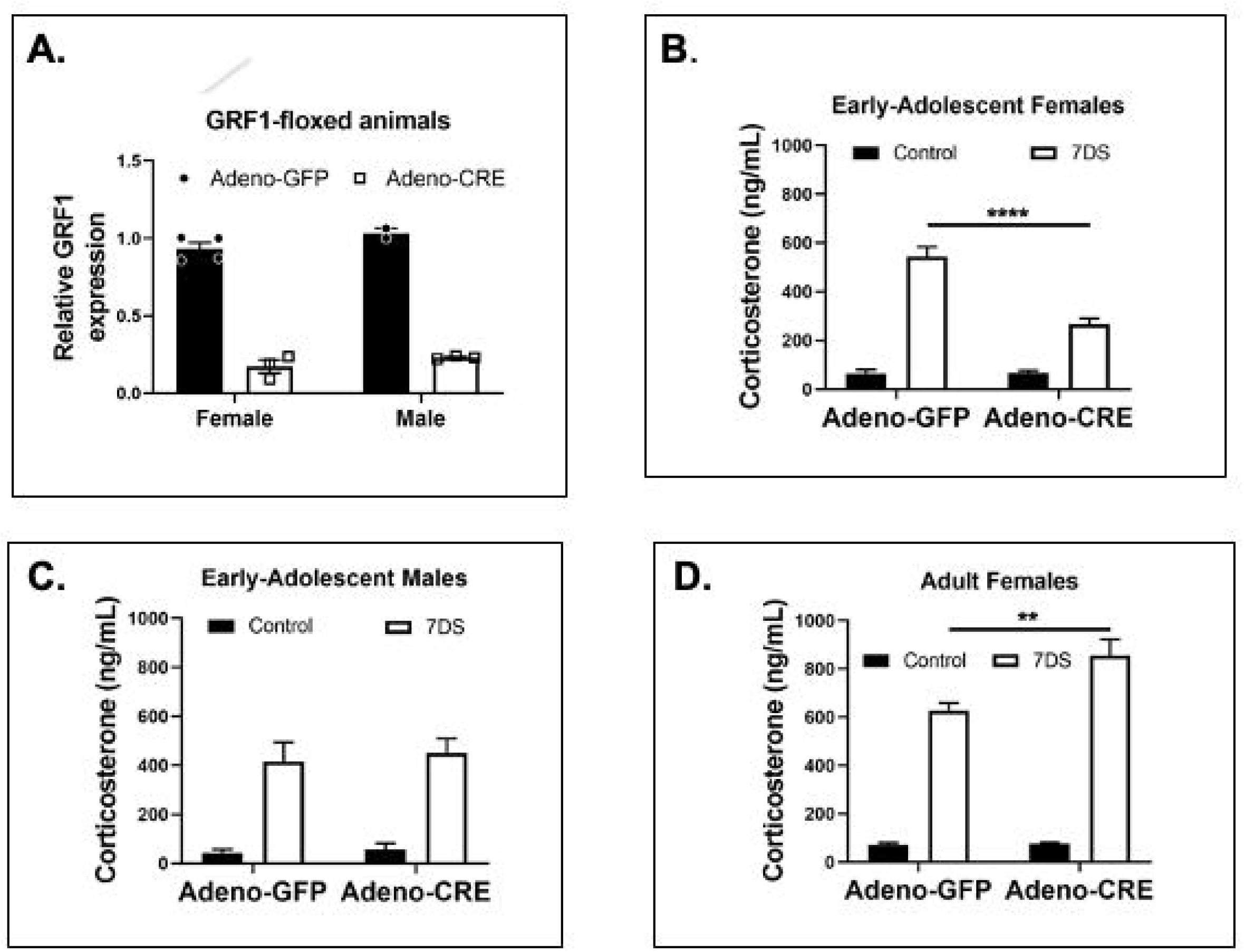
Inhibition of GRF1 function in the PVN of Early Adolescent (EA) female mice, but not in their male counterparts or adult female mice, suppresses HPA axis response to repeated restraint stress. **A.** Injection of adeno-CRE virus into the PVN of floxed EA females (Left) and floxed EA males (Right) effectively inhibits expression of GRF1 mRNA when compared to floxed animals injected with adeno-GFP as control. Each symbol represents an individual animal. B. Suppression of GRF1 expression in the PVN of EA females partially blocks elevated serum corticosterone in response to 7 exposures of restraint stress (30 min/day). Serum corticosterone levels measured after stereotactic injection of either Adeno-GFP or Adeno-CRE into the PVN of floxed GRF1 mice followed by control handling or 7DS (n=7 mice/group) **C.** The same as in **B.,** but for EA male mice (n=4 in each group) or **D.** Adult female mice (n=5 mice/group). Data presented as the mean +/− SEM. Analyses used: **A**) 2-way ANOVA, **B-D**) 3-way ANOVA. **P<0.01, ****P<0.0001.

After injection, baseline circulating CORT levels in unstressed animals were similar between Adeno-GFP and Adeno-CRE injected mice, consistent with our global GRF1 knockout mice (Uzturk et al. 2015) (Figs. 2B-D). However, after the final 30 min exposure to 7DRS, CORT levels were significantly reduced (~50%) in EA female mice with suppressed GRF1 expression when compared to their Adeno-GFP injected counterparts (Fig 2B). 3-way ANOVA (Factors: Sex/Age, CRE, 7DS) revealed a significant effect for Sex/Age (F_(2,52)_=30.06, *P*<0.0001) and 7DS (F_(1,52)_=488.7, *P*<0.0001), as well as the interaction between these two factors (F_(2,52)_=25.25, *P*<0.0001), the interaction between Sex/Age and adeno-CRE injection (F_(2,52)_=14.49, *P*<0.0001), and the interaction of all three factors (F_(2,52)_=14.31, *P*<0.0001), without a significant interaction for the adeno-CRE injection and 7DS factors (F_(1,52)_=0.0908, *P*=0.7644); Tukey’s multiple comparisons *post-hoc* tests revealed a significant decrease in CORT levels after 7DS between adeno-GFP and adeno-CRE injected early-adolescent females (q=8.947, *P*<0.0001). However, this suppression was not as complete as we found previously in global GRF1 knockout mice.

Interestingly, GRF1 knockdown reduced serum CORT levels in EA females to those similar to that found in control stressed EA male mice (see Fig. 2C bars on left), suggesting GRF1 plays a role in the enhanced sensitivity EA female mice display in response to 7DRS.

Importantly, GRF1 knockdown did not affect stress-induced CORT levels in EA male mice (Fig. 2C, bars on right). In addition, GRF1 knockdown did not suppress CORT level elevation in adult female mice (Fig. 2D). These findings are consistent with our previous work using global GRF1 knockout mice (Uzturk et al. 2015).

GRF1 knockdown actually led to a small increase in stress-induced CORT levels in adult females (Tukey’s multiple comparisons *post-hoc* tests revealed a significant increase in CORT levels after 7DS between adeno-GFP and adeno-CRE injected adult females (q=6.210, P=0.0029) (Fig 2D)). This raised the possibility of an indirect opposite effect of GRF1 knockdown in the other GRF1 expressing cell types in the PVN of adult females that express GRF1 (see Fig. 1A), as we don’t see the same effect when we knockdown GRF1 only in CRF cells (see below).

### Inhibition of GRF1 expression only in PVN CRF cells suppresses elevation of serum corticosterone levels induced by 7 days (30 min/day) of restraint stress (7DRS) specifically in early-adolescent (EA) female mice

Next, we investigated whether the suppression of elevated serum CORT levels in mice where GRF1 is knocked down only in the PVN after 7DRS (Fig. 1) is due, at least in part, to the loss of GRF1 in PVN CRF cells. Previous single cell RNAseq analysis demonstrated robust GRF1 expression in adult male PVN cells coexpressing CRF (Romanov et al. 2017) (also see http://linnarssonlab.org/hypothalamus/). First, we tested whether GRF1 is also expressed at the mRNA level in PVN CRF cells in EA females. To this end, GRF1 mRNA was quantified in brain slices from mice where only CRF cells express GFP. To tag CRF cells this way, the PVN of 21 day-old transgenic mice expressing CRE recombinase only in CRF cells (CRF-ires-CRE) (Taniguchi, et al. 2011) were injected with adeno-associated virus encoding a CRE-on construct that expresses GFP targeted to the cytoplasm by virtue of fusing it to the ribosomal protein L10a (see Materials and Methods and (Nectow, et al. 2017)), where GRF1 is expressed. Male and female EA as well as adult female mice where then exposed to 7 days of stress, when the sex and age dependent role for GRF1 is apparent. (Uzturk et al. 2015) (and see Fig. 1).

Because the L10a ribosomal subunit in GFP-L10 associates with actively transcribed mRNA, we could test whether EA female mice express GFP predominantly in CRF cells by quantifying CRF mRNA associated with GFP after its immunoprecipitation from cells, using the “TRAP protocol” (Nectow et al. 2017). As expected with successful immunoprecipitation, we found an enrichment of CRF mRNA of ~7-20 fold in precipitated mRNA compared to unbound mRNA, in RNA extracted from the PVN from infected mice. Then, we compared the ratio of GRF1 to CRF mRNA in the precipitated vs unbound fractions from both male and female EA mice and found that GRF1 mRNA was clearly enriched in the precipitated population of both mRNAs, although not to the level of CRF (Fig 3A).

**Figure. 3.**
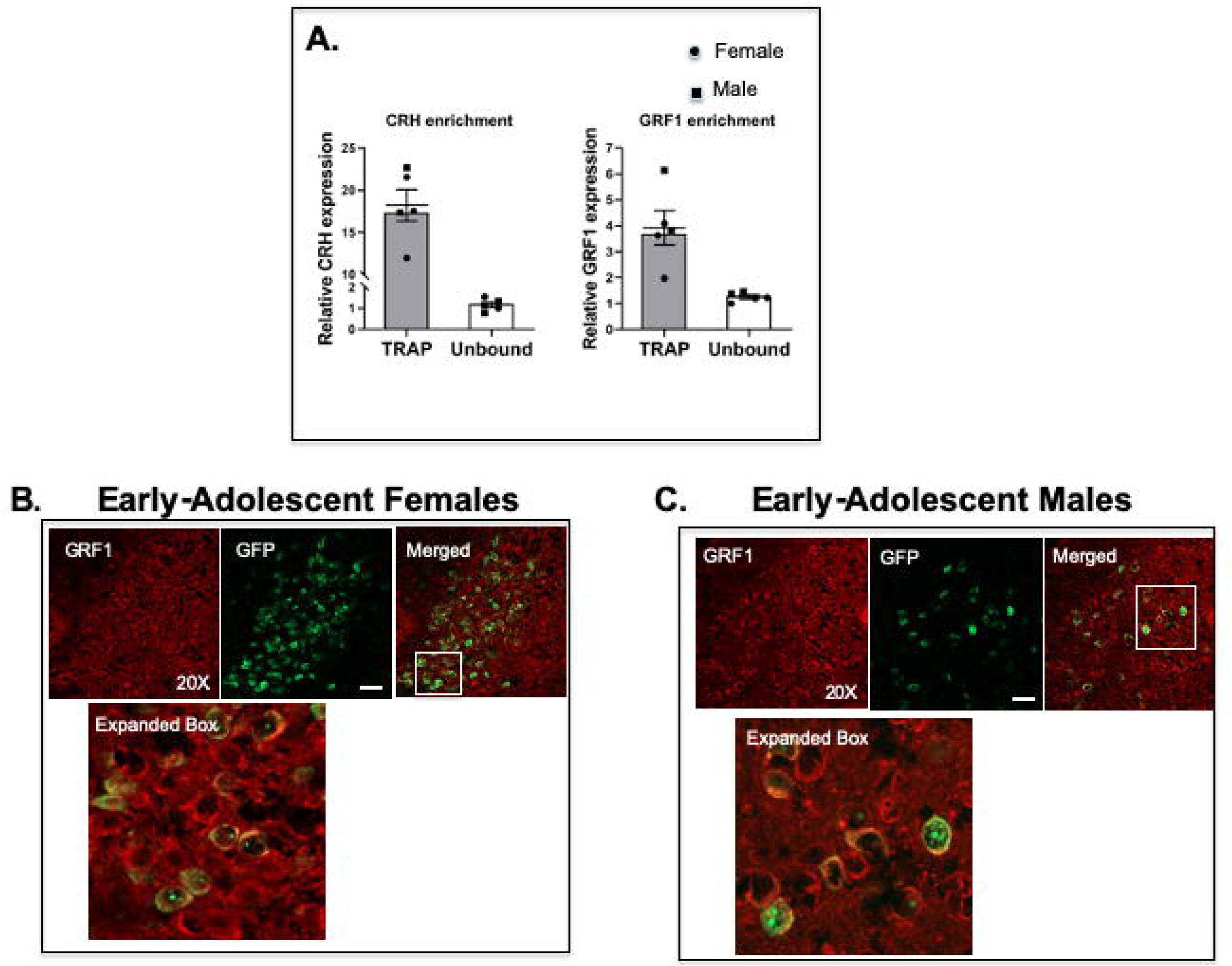
GRF1 is expressed in CRF cells of the PVN. The PVN of mice expressing CRE specifically in CRF cells were infected with a Lenti-virus expressing cytosol-targeted GFP-L10a fusion protein to allow isolation of ribosome associated mRNAs specifically in CRF cells as well as specifically label CRF cells with GFP. **A.** CRF cell mRNA was purified from total PVN mRNA by immunoprecipitation of GFP-L10a and compared with unbound non-selected mRNA for CRF (Left) and GRF1 (Right) mRNA levels detected by qPCR. Results from three EA females (circles) and two EA males (squares) are shown. Alternatively, brains sections were immunostained with anti-GRF1 antibody following 7DS for EA females (**B**), and EA Males (C) The insert bars for each set of images shows the percentage of CRF cells (GFP^+^ cells) that also stained positively for GRF1 expression (scale bar, 50 μm).

The finding that GRF1 is enriched in mRNA from CRF cells compared to total PVN mRNA, but not the extent that CRF is (only ~1/3 the level) was also expected. This is because, unlike CRF, GRF1 is expressed throughout the PVN (see Fig. 1). However, while GRF1 is expressed in excitatory neurons, it is not expressed in all cells such as glia, which make up a significant fraction of PVN cells (Stern and Filosa 2013).

To confirm that GRF1 is expressed in EA female CRF cells at the protein level, brain slices containing the PVN were stained with GRF1 antibody and the level of both red (GRF1) and GFP (green) were visualized. These images show representative examples of cells co-stained with GRF1 and CRF in both EA female and EA male CRF cells (Fig. 3B,C), despite the fact that GRF1 is not needed in the PVN of the latter mouse population for their full HPA axis response to repeated restraint stress (see Fig. 2).

Next, the consequence of inhibiting GRF1 only in CRF cells was investigated. A protocol similar to that described in Fig. 2 was employed, where GRF1 knockdown was targeted to the entire PVN of floxed GRF1 mice using a virus with a constitutive promoter driving CRE. However, here a lentivirus expressing CRE from a CRF cell-specific promoter (Lenti-CRF-CRE (Melon, et al. 2018)) was used in its place, or a virus expressing GFP as control. To confirm that this protocol actually reduced GRF1 levels specifically in CRF cells, wild-type or floxed GRF1 mice were co-injected with the CRF-CRE expressing lentivirus along with the GFP-L10 expressing virus used in Fig 3 to identify CRF expressing cells. Then, GRF1 antibody staining in CRF cells was compared in virus injected wild-type mice or floxed GRF1 mice. As expected, GRF1 staining in GFP-L10 expressing CRF cells was drastically reduced in virus-injected floxed GRF1 mice compared to wild-type mice (Fig 4A open bars). This level of reduction was comparable to what is seen in GRF1 staining reductions observed in animal wide GRF1 knockout mice (Fig. 4A, gradient bars, also see Fig. 1A and 1B). Finally, GRF1 knockdown was specific for CRF cells because there was no reduction in GRF1 staining seen in non-CRF expressing (non-GFP expressing) cells of the PVN in virus injected floxed GRF1 mice (Fig 4A, solid bars).

**Figure. 4.**
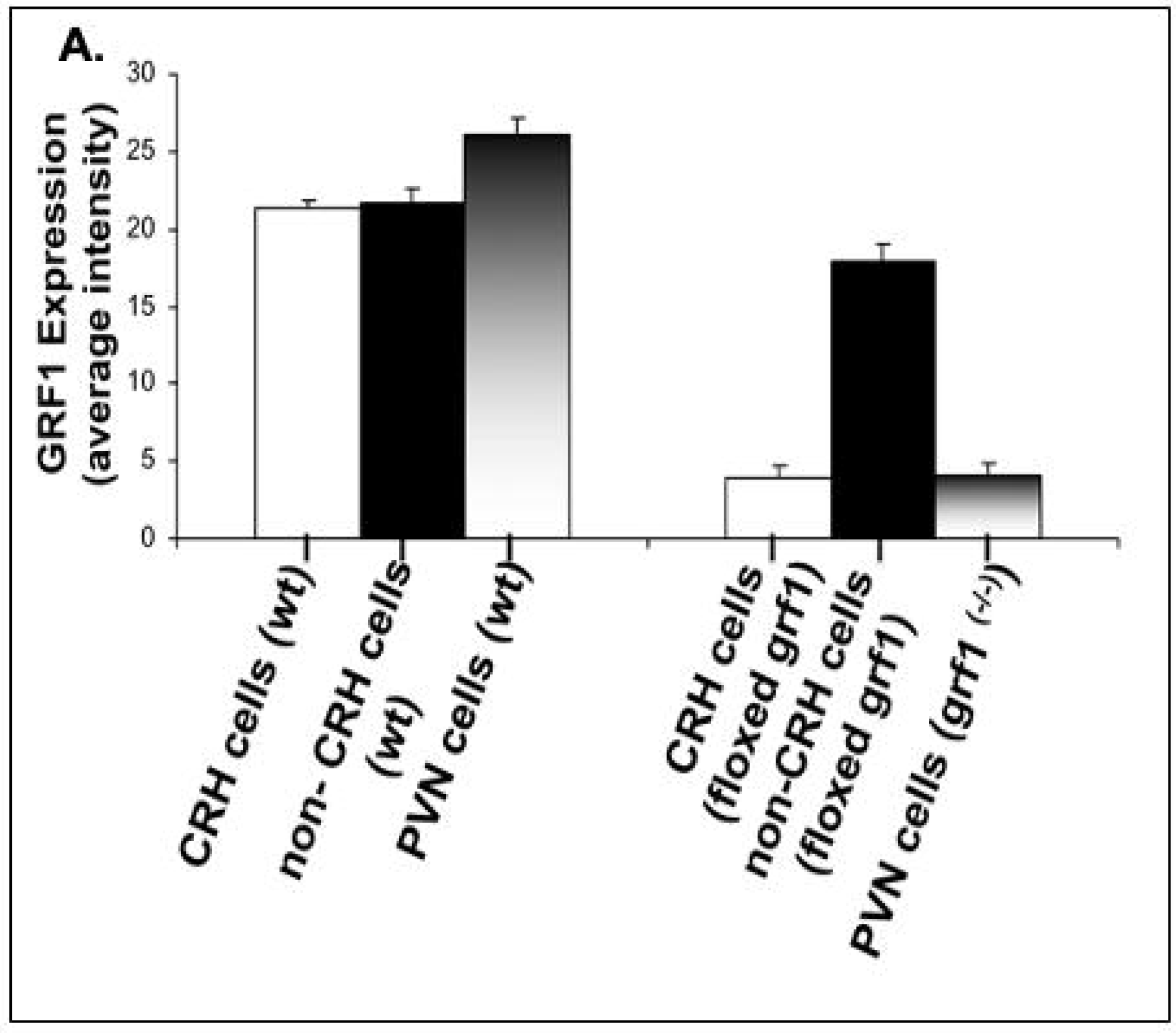
Demonstration of suppressed GRF1 expression specifically in CRH cells of the PVN. Levels of GRF1 protein expression in WT mice (Left) in CRF cells (white bar), non-CRF expressing cells (black bar), or the average cell in the PVN (gradient bar). The same expression was quantified in floxed mice (Right) after injection of lentivirus expressing CRE recombinase under the control of the CRF promoter again looking in CRF cells (white bar) or non_CRF expressing cells (black bar), compared to the average PVN cell in global GRF1^−/−^ animals (graded bar).

Similar to PVN injection of Adeno-CRE virus (Fig. 2), injection of CRF-CRE virus into EA female floxed GRF1 knockout mice led to a ~50% decrease in CORT response immediately after the 7^th^ exposures to restraint stress, compared to their GFP injected counterparts (Fig. 5A). 3-way ANOVA (Factors: Sex/Age, CRE, 7DS) revealed a significant effect for Sex/Age (F_(2,51)_=11-40, *P*<0.0001) and 7DS (F_(1,51)_=324.6, *P*<0.0001), as well as the interaction between these two factors (F_(2,51)_=11.17, *P*<0.0001), the interaction between Sex/Age and CRE (F_(2,51)_=4.480, *P*=0.0161), and the interaction of all three main factors (F_(2,51)_=4.632, *P*=0.0142), without a significant interaction between CRE and 7DS (F_(1,51)_=3.073, *P*=0.0856). Tukey’s multiple comparisons *post-hoc* tests revealed a significant decrease in CORT levels after 7DS between adeno-GFP and adeno-CRE injected early-adolescent females (q=6.920, *P*=0.0006).

**Figure. 5.**
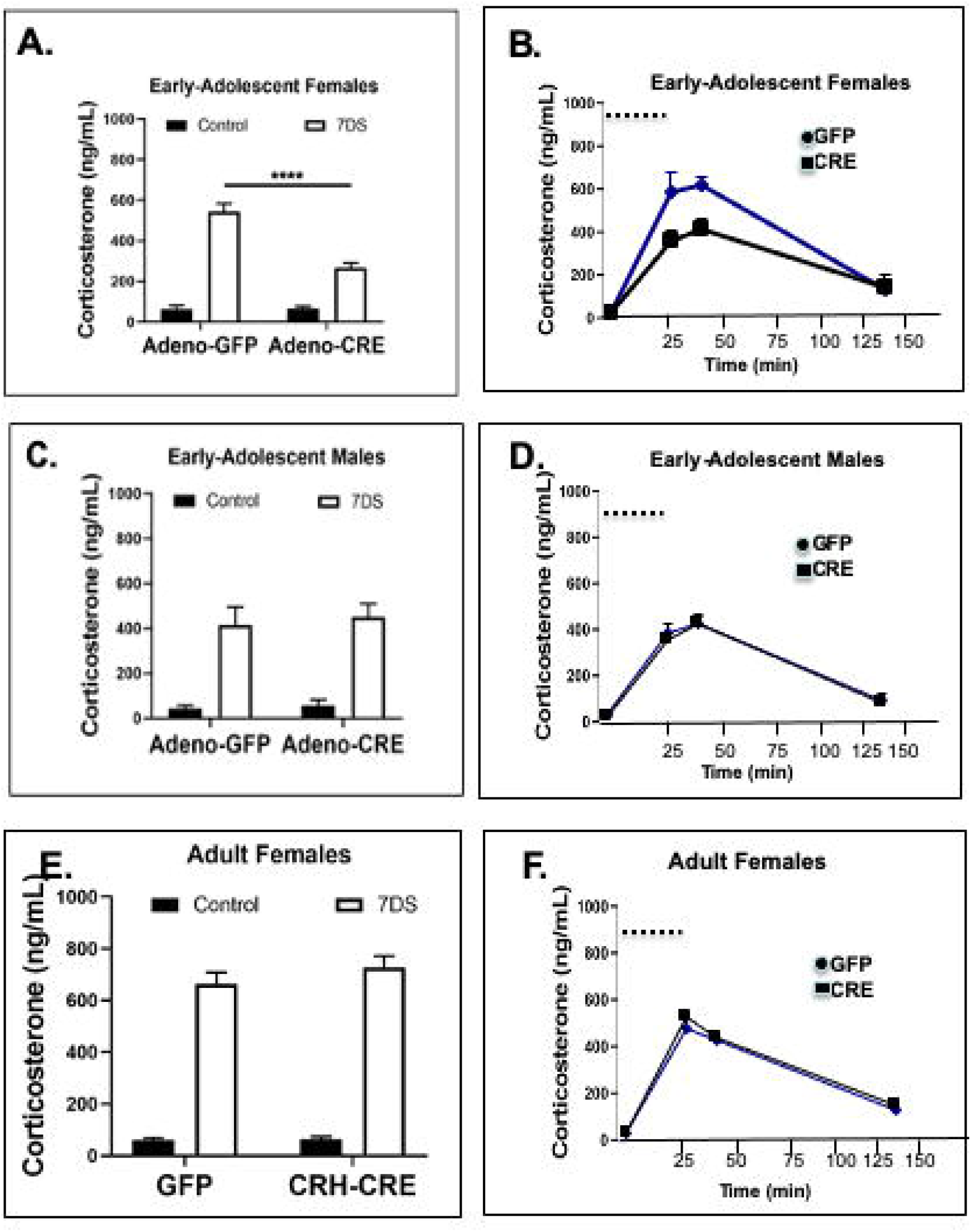
Knockdown of GRF1 expression specifically in CRF cells of the PVN suppresses elevated CORT levels in response to 7DRS in EA females, but not EA males or adult females. **A.** Suppression of GRF1 expression in the CRF cells of the PVN of EA females partially blocks elevated serum corticosterone response to 7 exposures of restraint stress (30 min/day). Serum corticosterone levels measured after stereotactic injection of either CRF-cre expressing lentivirus, or a GFP-expressing virus into the PVN of floxed GRF1 mice followed by control handling or 7DS. N=4 mice/group for both control groups and GFP-injected 7DS EA females, and n=7 mice for CRF-Cre injected 7DS EA females. **B.** Time course of serum CORT response. Data in **A** is included along with CORT levels measured 15 min and 90 minutes following the end of the stressor. N=3 mice/group C. Suppression of GRF1 expression in the CRF cells of the PVN of EA males does not affect serum CORT response to 7DRS N=7 mice/group **D.** Time course of serum CORT response. Data in **C** is included along with CORT levels measured 15 min and 90 minutes following the end of the stressor. N=3 mice/group E. Suppression of GRF1 expression in the CRF cells of the PVN of adult females does not affect serum CORT response to 7DRS N=7 mice/group **F.** Time course of serum CORT response. Data in **E** is included along with CORT levels measured 15 min and 90 minutes following the end of the stressor. N=3 mice/group. Data in **A, C, E** analyzed together by 3-way ANOVA, data in **B, D, F** analyzed separately using a Mixed Effects model. ***P<0.001

Moreover, performing a time course of serum CORT levels after the 7^th^ stress exposure showed a~50% reduction not only 30 min after beginning of the stress protocol but also 15 minutes later (Fig. 5B). Mixed effects model (Factors: Time, Cre-injection) for EA females revealed a significant effect for time (F_(1.319,8.792)_ = 46.48, *P*<0.0001) with a significant interaction between the two factors (F_(3,20)_ = 3.194, *P*=0.0457), the effect of CRE-injection was just outside of significance (F_(1,12)_=4.689, *P*=0.0512); Sidak’s *post-hoc* multiple comparisons revealed a significant difference between GFP and CRE injected animals immediately after the stress (t=4.831, *P*=0.0116), and 15 min after the stress (t=10.05, *P*=0.0384).. By 135 minutes, CORT levels in both control and GRF1 knockdown mice returned to almost normal levels and a significant difference was no longer observable. (Fig 5B). Sidak’s *post hoc* test did not reveal a significant difference in CORT levels after 7DS between GFP and Lenti-CRH-CRE injected animals (t=0.2156, *P*=0.9995).

Again, no significant effect was observed in EA males (Fig. 5 C and D) or adult females (Fig. 5E and F) immediately after the 7^th^ stress exposure as well as up to 120 min after initiation of stress. This occurred despite the fact that GRF1 is expressed at similar levels in EA male and EA female CRF cells (Fig. 3). Mixed effects model (Factors: Time, Cre-injection) for EA males revealed a significant effect for time (F_(1.585,9.508)_ = 64.50, *P*<0.0001) without a significant interaction between the two factors (F_(3, 18)_ = 0.0384, *P*=0.9896); Sidak’s *post hoc* multiple comparisons test did not reveal a significant difference between GFP and CRE injected animals before the 7^th^ restraint stress, immediately after the stress, 15 min after the stress, or 105 minutes after the stress. Mixed effects model (Factors: Time, Cre-injection) for adult females revealed a significant effect for time (F_(1.756,10.39)_ = 161.2, *P*<0.0001) without a significant interaction between the two factors (F_(3,28)_ = 0.3276, *P*=0.8054); Sidak’s *post hoc* multiple comparisons test did not reveal a significant difference between GFP and CRE injected adult female animals before the 7^th^ restraint stress, immediately after the stress, 15 min after the stress, or 105 minutes after the stress.

Thus, GRF1 in EA female CRF cells is required for a full CORT response of the PVN to 7 days of 30min/day restraint stress, but not their EA male or adult female counterparts.

Furthermore, as seen in total PVN knockdown (see Fig. 2), knockdown of GRF1 in CRF cells reduced the CORT response in EA females to levels near that for EA males, suggesting that GRF1 may be functioning in these cells to account for the increased expression of CORT following stress seen in EA females when compared to their male counterparts (Tukey’s multiple comparisons *post-hoc* tests revealed a significant difference in CORT levels after 7DS between EA males and EA females injected with adeno-GFP (q=6.327, P=0.0023) (Fig 5A, C)).

### Inhibition of GRF1 expression only in PVN CRF cells suppresses the anxiolytic effect of 7 days (30 min/day) of restraint stress in early-adolescent (EA) female mice

*We* showed previously that the paradoxical decrease in anxiety observed previously after exposure of adolescent rodents to other types of stressors (Toledo-Rodriguez and Sandi 2011) also occurs in B16 mice exposed to restraint stress (Uzturk et al. 2015). In particular, we showed that 24 hrs after 7 days of restraint stress, EA females spend more time than control animals in the open arms of the elevated plus maze. Interestingly, we found that this was not true for their EA B16 male mice counterparts. Moreover, this phenotype was blocked in EA female GRF1 knockout mice, consistent with the block in CORT response these mice display (Uzturk et al. 2015). Thus, we exposed EA female mice whose GRF1 expression was inhibited specifically in PVN CRF cells to this test. Even though the CORT inhibition following 7DRS was not complete in these mice, (See Fig. 4B.), anxiolytic behaviors on the EPM were completely suppressed 24 hrs after cessation of the stressors (Fig. 6C). 2-way ANOVA (Factors: 7DRS, CRE-injection) revealed an overall effect for 7DS (F(1,12)= 5.061, *P*=0.0440) without a significant interaction between factors (F(1,12)=2.558, *P*=0.1357). Sidak’s *post hoc* multiple comparisons test revealed a significant difference following 7DS only in GFP injected animals (t=2.722, *P*=0.0367).

**Figure. 6.**
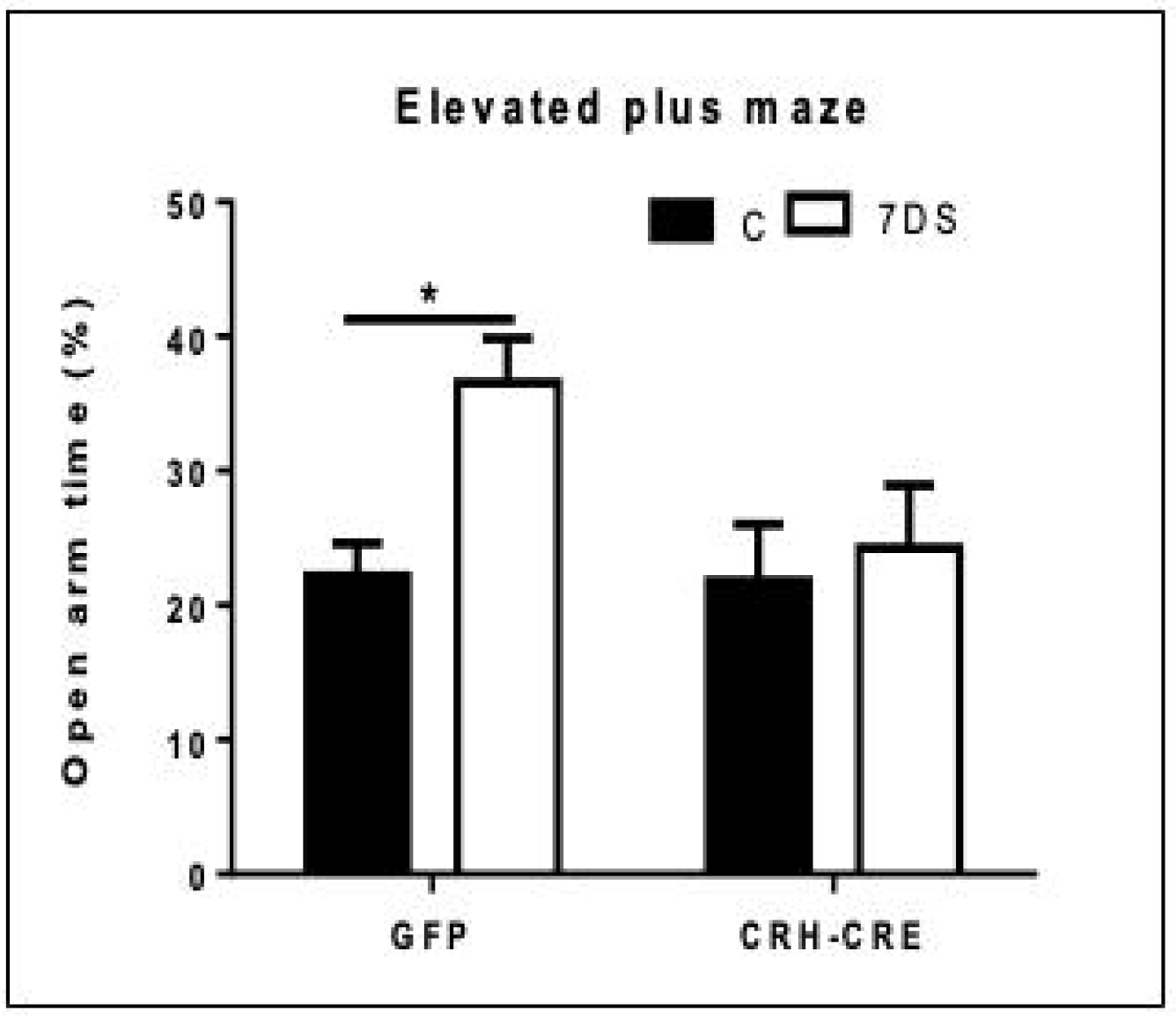
Knockdown of GRF1 expression specifically in CRF cells of the PVN suppresses anxiolytic effects of 7DRS. Following stereotactic injection of either CRF-cre expressing lentivirus or a GFP-expressing virus into the PVN of floxed GRF1 mice, animals were either exposed to 7DRS or control handling. On the following day, they were assayed for time spent in the open arms of the elevated plus maze (n=4 mice/group). Data analyzed by 2-way ANOVA. \**P*<0.05.

### Knockdown of GRF1 in CRF cells in the PVN hypothalamus does not influence the HPA axis response to short-term restraint stress

We showed previously that the loss of GRF1 specifically in the CA1 hippocampus of early-adolescent GRF1 knockout mice is responsible for the super elevated levels of circulating CORT levels observed in them after short-term stress (30 min/day of restraint for 3 days)(Uzturk et al. 2015). To explain how this can occur if GRF1 is needed in CRF cells to generate a full CORT response, GRF1 was knocked down specifically in CRF cells (as in Fig. 4B,C,D), and mice were then exposed to only 3 days of restraint stress at the end of the experiment. In this case, the Lenti-CRF-CRE virus was also injected into the PVN at pn 21, but restraint stress was initiated 11 rather than 7 days later, so that the mice were exposed to the virus for the same amount of time and the mice were the same age at the end of the experiment as those from 7 days of restraint stress. In contrast to the inhibitory effect observed from similarly injected EA female mice exposed to 7 days of stress (see Fig. 4A), no effect was observed right after 3 days of restraint stress (Fig. 7A). (2-way ANOVA (Factors: CRE, 3DS) revealed a significant effect for 3DS (F_(1,12)_=190.3, *P*<0.0001) without a significant interaction between factors (F_(1,12)_=0.6530, *P*=0.4348). Sidak’s *post hoc* multiple comparisons test did not reveal a significant difference between GFP and CRE-injected EA females following 3DS. Additionally, there was no significant difference in CORT levels after 3DS between GFP and Lenti-CRF-CRE injected animals 15 minutes following completion of the stressor, or 105 minutes following completion of the stressor (Fig. 7B). Mixed effects model (Factors: Time, Cre-injection) for EA females following 3DS revealed a significant effect for time (F_(1.346,8.977)_ = 130.5, *P*<0.0001) without a significant interaction between the two factors (F_(3,20)_ = 0.3149, *P*=0.8144); Sidak’s *post hoc* multiple comparisons test did not reveal a significant difference between GFP and CRE injected EA female animals before the 3^rd^ restraint stress, immediately after the stress, 15 min after the stress, or 105 minutes after the stress.

**Figure 7.**
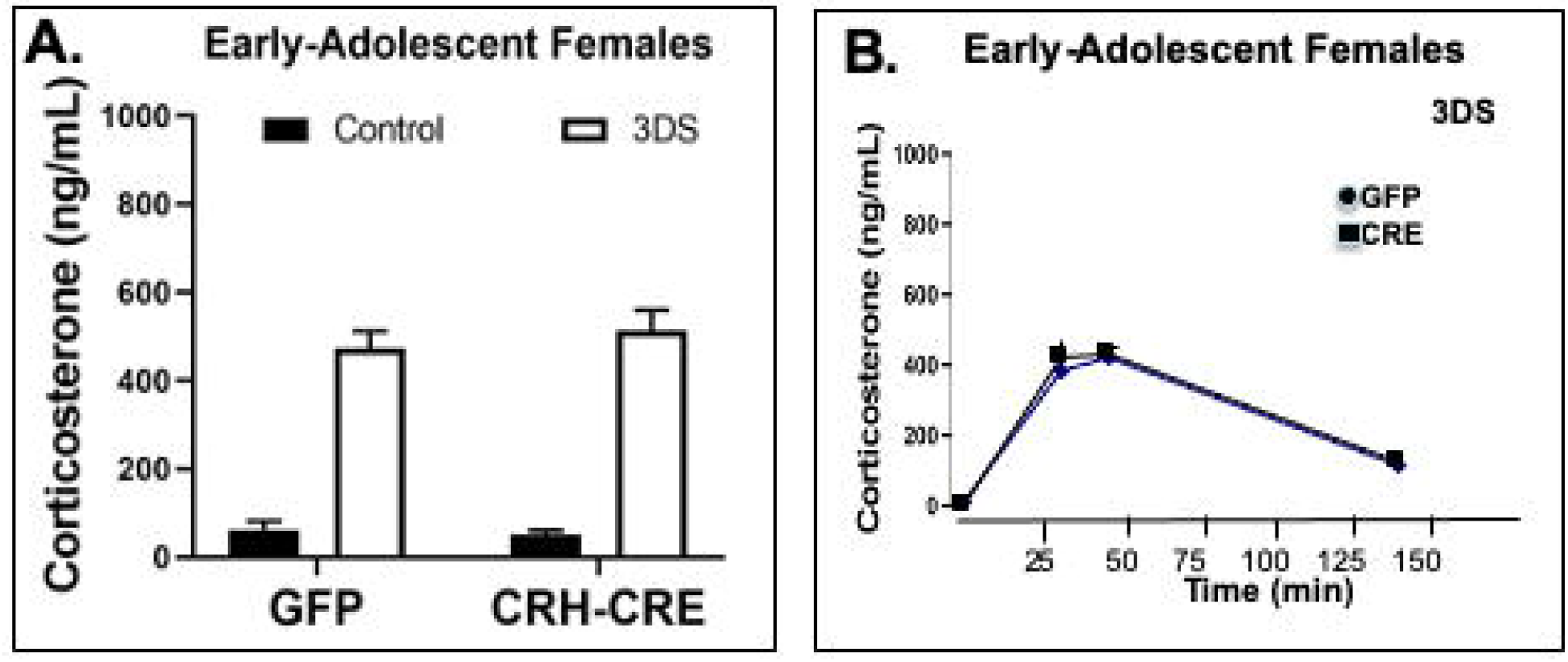
Knockdown of GRF1 expression specifically in CRF cells of the PVN of EA females does not suppress serum CORT response to 3 days of exposure to restraint stress. After stereotactic injection of either a GFP-expressing or CRF-cre expressing lentivirus into the PVN of floxed GRF1 mice, they were either exposed to 7DRS or control handling. Then serum CORT levels were measured A. right after the end of the last stress exposure; **B.** Time course of serum CORT response. Data in **A** is included along with CORT levels measured 15 min and 90 minutes following the end of the stressor. N=3 mice/group. Data in **A** analyzed by 2-way ANOVA, and in **B** analyzed by Mixed Effects model.

These findings show that GRF1 begins to contribute to HPA axis regulation in CRF cells in EA females only after additional repeated stress exposures. This implicates GRF1 in a mechanism of PVN CRF cell plasticity that may be specific to EA females, since it is not found in their EA male or adult female counterparts.

## DISCUSSION

The findings reported here show that GRF1 contributes to the HPA axis response to repeated restraint stress (7 days of 30 min/day) in EA females, at least in part, via its function in PVN CRF cells of the hypothalamus. We show that it not only contributes to HPA axis regulated serum CORT levels immediately and 15 minutes after the last stress exposure, but also decreased anxiety detected 24 hours later. In contrast, GRF1 in CRF cells does not contribute to HPA axis regulation of CORT in EA male or adult female mice. Thus, these results have implications for both sex and age-dependent differences in the stress response. They are also relevant to the process of CRF cell plasticity associated with chronic stress in EA females, since we showed that GRF1 in CRF cells of this population does not contribute to the PVN hypothalamus response to shorter-term restraint stress (3 days of 30 min/day). Finally, knocking down GRF1 in CRF cells only partially blocked the serum CORT response to stress while global knockdown of GRF1 led to an almost complete block of this response(Uzturk et al. 2015), GRF1 in other brain regions, possibly in other CRF secreting cell types, may also contribute to HPA axis regulation.

The idea that CRF cells in EA females are distinct from their EA male and adult female counterparts is not surprising since both adolescents and females are known to respond to stress differently from males and adults (Bale and Epperson 2015). In particular, women are more likely than men to suffer from depression (Bangasser and Valentino 2014) and PTSD (Bremner, et al. 1997; Nemeroff, et al. 1984) and respond better to a certain class of antidepressants than others (Kornstein, et al. 2000). Sex-dependent properties of CRF cells may be involved since isolating preadolescent female, but not male, mice from littermates for <24 hr increased first spike latency and decreased excitability of CRF neurons (Senst et al. 2016).

In general, females and adolescents tend to generate enhanced CORT responses to stress. We observed sex dependence here comparing male and female EA mice after 7 exposures to restraint stress (compare Figs. 2B and C and 4B and C). Interestingly, GRF1 knockdown in CRF cells of EA females reduced their CORT response to levels similar to those observed in EA males, implying that GRF1 function specifically in EA females contributes to the enhanced HPA axis observed in these animals.

Interestingly, adolescent rats exposed to psychogenic stress, involving 3 days of exposure to a variety of stressors, display what appears to be a paradoxical effect of decreased anxiety associated with increased risk taking (Toledo-Rodriguez and Sandi 2011). We previously observed a similar anxiolytic effect, increased time in the open arms of the EPM after exposure of EA female B16 mice to the repeated restraint stress paradigm studied here. In contrast to studies in rats, we did not observe this effect in EA males (Uzturk et al. 2015). Consistent with inhibition of elevated serum CORT levels in EA female animal wide GRF1 knockout mice, this anxiolytic effect of repeated restraint stress was also completely blocked in these mice. Here, we also observed a complete blockade of the anxiolytic effect of this stressor when GRF1 expression was blocked only in CRF cells, even though it only partially blocked stress-induced elevation of CORT levels to those found in males. Thus, it is possible that GRF1 in CRF cells plays a role in stress-induced risk taking, a behavior linked to dangerous outcomes, specifically in adolescent females. Finally, that GRF1 is present in EA male CRF cells implies it plays a different role in them that remains to be determined.

EA in mice (pn 21–35), the period where GRF1 functions to control the HPA axis in female CRF cells, is thought to be similar to human ages of 10–14 (Burke and Miczek 2014). It is a period of rapid brain development and high vulnerability to excessive stress (Andersen, et al. 2008). Risk of depression upon early stress experiences was shown to be highest around early-adolescence and decreases as humans age through mid- and late-adolescence (Andersen and Teicher 2008). Moreover, EA girls are more prone than their male counterparts to anxiety, depression and eating disorders, as well as suicidal ideation (Hankin et al. 1998; Skogli et al. 2013). Thus, dysregulation of GRF1 function could contribute to these age-related maladies.

Some of these studies have demonstrated that age and sex differences in the HPA axis can arise from the influence of gonadal hormones in adulthood or post pubertal periods (Handa and Weiser 2014). However, we showed that GRF1 participates in HPA axis function before puberty and ovariectomy has no effect on this role (Uzturk et al. 2015). Thus, in this case, the sex bias of GRF1 function likely results from perinatal testosterone surges in males that makes them independent of GRF1 or from other sex chromosomal effects, which are much less well studied in the context of the stress response (Arnold 2009). We cannot discount the possibility that specific GRF1 function is also present in females younger than EA because we have avoided restraining mice before weaning.

Another key finding of this study is that although knockdown of GRF1 function in CRF cells suppresses HPA axis function after 7 exposures to restraint stress, it has no effect after 3 such exposures. This finding explains how we could have found enhanced CORT response after 3 restraint stress exposures in our previous animal wide GRF1 knockout studies (Uzturk et al. 2015), since we now know GRF1 is only needed in CRF cells after additional stress exposures. This implies that the requirement for GRF1 in CRF cells for a full HPA axis response to stress only after 7 exposures is part of a form of CRF cell plasticity that is distinct in EA females. CRF cell plasticity has been well documented at the level of altered cell excitability (Bains, et al. 2015; Maguire 2014) after various stress paradigms, but sex- and agespecific forms have not. Disturbed plasticity is likely involved in some forms of stress-related disorders, and this form that incorporates GRF1 function could be particularly relevant to those that arise in young women (Bains et al. 2015; Maguire 2014),

That GRF1 function would be an important feature of CRF cells only before puberty in females, and only after repeated stress was surprising because we showed previously that GRF1 contributes to synaptic plasticity, learning, and memory in both sexes, as well as in both adolescents and adults (Jin et al. 2013; Li et al. 2006). Moreover, we documented here that GRF1 is present at similar levels in CRF cells from stressed EA females, and EA males, even though it is only needed for a full HPA axis response in the former. Interestingly, we have observed in a previous study that although GRF1 is present in the hippocampus of 1-month old mice, it does not begin to contribute to the process of LTP induced by calcium permeable AMPA receptors and contextual discrimination until mice reach 2 months of age (Jin et al. 2013). Thus, what remains to be discovered is not only which GRF1-mediated signaling cascades participate in these CRF cells to generate proper HPA axis control, but also what is different about repeatedly stressed EA female CRF cells that GRF1 is required in them, but not their EA male or adult female counterparts.

Plasticity of CRF cells to repeated stress has been shown to involve changes in excitability that makes the system more sensitive via stress-induced changes in excitatory/inhibitory input ratio (Bains et al. 2015; Maguire 2014). GRF1 mediates the actions of both excitatory NR2B type NMDA and calcium-permeable type AMPA glutamate receptors involved in synaptic plasticity regulation in the hippocampus. Thus, GRF1 could regulate upstream activation of CRF cells because these cells are also stimulated by glutamate. This mechanism would suggest that a reason for a distinct GRF1-mediated regulation pathway in EA females is to allow them to respond to a distinct set of environmental cues specific to EA females.

Alternatively, GRF1 could function downstream of cell activation by participating in enhanced CRF gene expression, which is known to occur after stress. Finally, GRF1 could function in excitation/CRF secretion coupling. For example, a downstream function of GRF1 is to activate the Ras/Erk Map kinase pathway, which is known to promote neurotransmitter secretion through phosphorylation of synaptic regulatory proteins (Chi, et al. 2003). This mechanism would imply that the purpose of a distinct GRF1 signaling pathway in EA females is to generate a unique HPA axis response required for this population of animals.

Of course, all of these potential functions for GRF1 in CRF cells would have to be activated in EA females, but not in comparable males or adult females, and only after 7 exposures to restraint stress. That means that a unique feature(s) of CRFs cells in repeatedly-stressed early-adolescent females might be to engage an otherwise inactive GRF1, or alter its signaling specificity to bring about a new function. In fact, precedent exists for sex-dependent specificity differences in neuroendocrine signaling cascades. For example, females display a decreased ability of the CRF receptor to associate with β-arrestin 2, which biases signaling through Gs-related vs arrestin-related signaling pathways (Valentino, et al. 2013).

## ACKNOWLEDGEMENTS

Authors have no conflicts of interest. We thank Eric Schmitt, Rockefeller Univ for CRE-inducible”FLEX” L10a-GFP Adeno-associated virus and anti-GFP antibody, and Dong Kong, Tufts University School of Medicine, for the CRF-ires-Cre mice. This work was supported by funds to LAF from NIH # R01MH107536.

There are no conflicts of interest.

## AUTHOR CONTRIBUTIONS

Shan-xue Jin and David Dickson helped design and carry out all experiments. Jamie Maguire and Larry Feig helped design experiments. David Dickson and Larry Feig wrote the manuscript.

## MATERIALS AND METHODS

### Animals Care Facilities

All mice included in this study were housed in temperature, humidity, and light-controlled (14 hour on/10 hour off LD cycle) rooms in a fully staffed dedicated animal core facility led by on-call veterinarians at all hours. Food and water were provided ad libitum. All procedures and protocols involving these mice were conducted in accordance with and approved by the Institutional Animal Care and Use Committee of the Tufts University School of Medicine, Boston MA. CRF-ires-CRE mice were obtained from Dong Kong, Tufts University School of Medicine via Jackson Labs (Stock No.: 012704) and bred in-house in a C57-BL6 background (https://www.jax.org/strain/012704).

#### Statistical Methods

All statistical analyses were performed in GraphPad Prism v.8.3.1. All grouped analyses comparing CORT measurements with/without stress, with GFP or CRE injection, and including EA male, EA female, and adult female mice were analyzed using a 3-way ANOVA (main effects: Age/Sex [Three-level factor: EA male, EA female, and adult female], 7DS, CRE), and F-statistics and p-values are reported in the text for all significant main effects and all interactions. For analysis of the knockdown of GRF1 mRNA in adeno-CRE injected floxed animals, as well as the CRF-Cre injected EA female mice after 3DS, analysis was carried out using a 2-way ANOVA (Factors: Cre, 3DS) in the same fashion as described above. For time course studies, analyses were carried out using Mixed Effect models for each Age/Sex of mouse individually, as each time point was gathered from a distinct cohort of mice. *Post hoc* analyses comparing CORT measurements in the 7DS or 3DS condition between GFP and CRE injected animals were corrected for multiple comparisons using Sidak’s method, in the case of 2-way ANOVAs to assess differences in CORT following 7DS or 3DS within groups, for the EPM analysis, and for Mixed Effect models comparing CORT levels at different time points. Tukey’s method was used in the case of 3-way ANOVAs where all means are compared to all other group means. All *post hoc* analyses were reported with the t-statistic and the multiplicity adjusted p-value. For all analyses, a P<0.05 was considered significant.

#### Viruses

AAV-CRE as obtained from Virovek; Lenti-CRF-CRE was obtained from the Virology Core of Emory University; CRE-inducible L10a-GFP Adenovirus was obtained from Eric Schmidt (Rockefeller University)

### Generation of floxed-GRF1 mice

The conditional knockout of GRF1 was accomplished by conditional deletion of a 96 bp coding region in the DH domain in exon 6, and the insertion of loxP sites flanking exon 6. The FRT-flanked selection cassette was removed in vivo by crossing with recombinase deleter mice (see Suppl. Fig. 1). Recombinant mice were back-crossed 6 times into the C57-B16 background.

### RNA extraction

For RNA extractions from whole PVN, total RNA was isolated using the Single Cell RNA Purification Kit from Norgen Biotek Co. according to the manufacturers protocol. All RNA isolates were assessed for concentration and purity using a Nanodrop 2000c Spectrophotometer (Thermofisher Scientific) and/or an Agilent 2100 Bioanalyzer.

### Real-time qPCR

Relative gene expression for all samples was determined using the iScript cDNA synthesis kit, and SYBR Green MasterMix purchased from Thermofisher Scientific. 75 ng of total RNA from whole PVN extractions were used in the initial cDNA synthesis step. Real-Time PCR was performed for each target and sample in triplicate on a StepOnePlus PCR System (Applied Biosystems). All data was analyzed using the Comparative ΔΔCT method to generate relative expression data using *rps29* as the internal loading control. Primers used: *rps29* Fwd: 5’-GTCTGATCCGCAAATACGGG-3’, *rps29* Rev: 5’-AGCCTATGTCCTTCGCGTACT-3’; *rasgrf1* Fwd: 5’-ATCACCTCCTCCATCAACCG-3’, *rasgrf1* Rev: 5’-CTGCCATCTGATGACACAAGC-3’, *CRF* Fwd: 5’-GGCATCCTGAGAGAAGTCCCTC-3’, *CRF* Rev: 5’-ACAGAGCCACCAGCAGCATG-3’.

### Translating Ribosome Affinity Purification

Ribosome-bound RNA isolation was carried out with an optimized form of the TRAP protocol (Nectow et al. 2017) Briefly, we began by covalently binding anti-GFP antibodies (19C8,19F7, gift from Eric Schmitt, Rockefeller Univ) to magnetic Dynabeads (Invitrogen), and treating with IgG-free BSA to reduce background binding. Whole PVNs were homogenized using plastic pestles, spun down to remove cellular debris, and added to the anti-GFP beads for binding for 2 hours at 4° C. The unbound fraction for each sample was removed and kept on ice for later RNA extraction. The beads were then washed twice with a low strength KCl buffer, and 3 times with a high strength KCl buffer. RNA was then extracted from both the washed beads and the unbound fractions using the Single Cell RNA purification kit (Norgen), with slight modifications to improve yield. For the bead samples, 100 uL of lysis buffer was added to the beads before vortexing 5 times for 5s each, and letting rest at room temperature for at least 5 minutes. The beads were removed by magnet, and the extraction then proceeded normally. For the unbound samples, 1 volume (the total amount of unbound sample, typically ~150 uL) of lysis buffer was added, the sample was vortexed, and then 2 volumes of room temperature 70% EtOH were added before proceeding with the extraction normally. All samples were eluted in 20 uL of DEPC-treated H_2_O before quantification and quality assessment as described in the RNA extraction section above.

### Immunohistochemistry

Fourteen days after the stereotactic surgeries, mice were deeply anesthetized with ketamine/xylazine and transcardially perfused with 0.1 M phosphate buffered saline (PBS) followed by 4% paraformaldehyde (PFA 4%) dissolved in PB 0.1 M. Brains were extracted and post-fixed in PFA 4% for 24 h. Brains were transferred to 30% sucrose for 48–72 h before slicing 30 μm coronal sections through the extent of the PVN using a cryostat. Sections were stored in cryoprotectant at −20 °C until use. Each immuno-histochemical analysis was conducted from 30 μm sections spanning the PVN.

Free-floating sections were rinsed extensively in PBS with 0.3% Triton X-100 (PBS-T). Sections were blocked for 1 h at room temperature in PBS-T with 5% normal goat serum. Primary antibody, antiGRF1 (C 18) (Santa Cruz Biotechnology, Santa Cruz, CA, USA), was diluted in the blocking solution (1:200), incubated overnight at 4 °C, and rinsed three times for 15 min in PBS-T. The sections were then incubated for 1.5 h at RT with a Alexa 488 or Cy3-conjugated secondary antibody (1:300; Invitrogen). The stained sections were examined with a confocal microscope and images were captured with a CCD spot camera. Images were then analyzed using ImageJ. For CRF cell-specific GRF1 expression, CRF cells were first identified by their robust GFP expression throughout the cytoplasm, and were individually traced with an overlay (10-20 GFP^+^ cells per slice, at least 3 slices per animal). The overlay was then added to the GRF1 channel of the image to quantify the total fluorescence signal for GRF1 inside each of the outlined cells, calculated as Corrected Total Cell Fluorescence ([Total fluorescence of the cell] – ([Area of the cell]*background signal)) using the average of 3 background measurements for each image. Cells were said to be positive for GRF1 if the total fluorescence counted for the cell was at least double that of the background staining, given that GRF1 is robustly expressed in many cell types in the brain.

### Stereotactic Injections

Twenty-one or twenty-two-day-old mice were injected as previously described (Darcy, et al. 2013). Each injection consisted of 1 μl of adenovirus or lentivirus infused at a rate of 0.06 ml/min. An infusion pump controlling the plunger on the Hamilton syringe precisely regulated the rate of injection. The needle was then left in place for 8 min prior to withdrawal from the brain. 4-7 mice were used for each experiment. Half of floxed siblings were injected with CRE expressing virus for experimental samples and half with GFP expressing virus as controls.

### Restraint Stress

Restraint stress experiment were carried out as previously described (Uzturk et al. 2015). Briefly, mice were placed into custom-fit cone-shaped plastic bags and secured in place for 30 min every day. The cone has an open end so the mouse can breathe, but can otherwise not move for the duration of the stressor. Mice are always stressed between 12:00PM and 2:00PM to control for the circadian fluctuations of corticosterone production.

### Corticosterone (CORT) ELISA

CORT measurements were carried out as previously described using the Corticosterone ELISA kit from Enzo Life Sciences (Uzturk et al. 2015). We noticed that EA female mice used here (floxed-BL6 mice) display a greater CORT response that we found in B16 mice three years ago, which could be caused by different housing conditions over the years or a difference in the strain.

